# Perceptual phenotypes: Perceptual gains and losses in synesthesia and schizophrenia

**DOI:** 10.1101/443846

**Authors:** Tessa M. van Leeuwen, Andreas Sauer, Anna-Maria Jurjut, Michael Wibral, Peter J. Uhlhaas, Wolf Singer, Lucia Melloni

## Abstract

Individual differences in perception are widespread. Considering inter-individual variability, synesthetes experience stable additional sensations; schizophrenia patients suffer perceptual deficits in e.g. perceptual organization (alongside hallucinations and delusions). Is there a unifying principle explaining inter-individual variability in perception? There is good reason to believe perceptual experience results from inferential processes whereby sensory evidence is weighted by prior knowledge about the world. Different perceptual phenotypes may result from different precision weighting of sensory evidence and prior knowledge. We tested this hypothesis by comparing visibility thresholds in a perceptual hysteresis task across medicated schizophrenia patients, synesthetes, and controls. Participants rated the subjective visibility of stimuli embedded in noise while we parametrically manipulated the availability of sensory evidence. Additionally, precise long-term priors in synesthetes were leveraged by presenting either synesthesia-inducing or neutral stimuli. Schizophrenia patients showed increased visibility thresholds, consistent with overreliance on sensory evidence. In contrast, synesthetes exhibited lowered thresholds exclusively for synesthesia-inducing stimuli suggesting high-precision long-term priors. Additionally, in both synesthetes and schizophrenia patients explicit, short-term priors – introduced during the hysteresis experiment – lowered thresholds but did not normalize perception. Our results imply that distinct perceptual phenotypes might result from differences in the precision afforded to prior beliefs and sensory evidence, respectively.

## Introduction

Schizophrenia is a neuropsychiatric disorder that is not only characterized by positive symptoms like hallucinations and delusions, and disorganized thought, but also by perceptual *deficits* such as impaired perceptual grouping^1^, multisensory integration^2^, object recognition deficits^3-5^, and impaired low-level visual processing (e.g.^6,7^). Another condition is synesthesia, a form of altered perception in which specific stimuli (e.g., letters) consistently and automatically trigger vivid *additional* conscious experiences (e.g., color), described as being percept-like^8^. Synesthetic experiences are associated with activation of cortical areas (e.g., color-sensitive) known to process stimulus qualities of the synesthetic concurrent experience^9,10^. Synesthetes are aware their synesthetic experiences are not real, contrary to hallucinations or delusions in schizophrenia. Perception can be altered in synesthetes^11^. Reports for low-level visual processing are limited, including both enhanced and reduced sensitivity^12,13^. We investigated schizophrenia patients and synesthetes because for low-level, degraded visual stimuli involving letters, their perceptual experiences differ despite receiving the same input – impaired perception in schizophrenia patients and additional color experiences for grapheme-color synesthetes.

There is good reason to believe that perceptual experience results from inferential processes whereby sensory evidence is weighted by prior knowledge about the world^14-16^. Therefore, it is possible that different perceptual phenotypes stem from dissimilar weighting of sensory evidence and prior knowledge. Here, we investigated whether inter-individual differences in how top-down priors are balanced against sensory evidence during perceptual inference may explain the differences in perceptual experience of synesthetes and schizophrenia patients. Research implementing perceptual inference accounts of psychiatric disorders has increased substantially in recent years^17,18,19^. An emerging hypothesis is a failure of *precision weighting. Precision* of prediction errors (unexplained input) determines the strength or weighting (inverse of variance) of sensory evidence in relation to prior beliefs: high precision of sensory prediction errors refers to the excitability of neurons signaling new information^20,21^ and biases perception towards sensory evidence, reducing the influence of prior beliefs^22^. Similarly, precise top-down predictions bias perception towards prior beliefs^18,23^.

Here, we focus on schizophrenia and synesthesia and specifically investigate whether their diverse *perceptual phenotypes* can be explained by aberrant precision weighting in perceptual inference. In schizophrenia, high-precision prediction errors for sensory input are hypothesized, causing an overreliance on sensory evidence during sensory processing with reduced influence of top-down information^19,24^. This explains, the paradoxical resistance to perceptual illusions observed in schizophrenia^5,25^. For example, while healthy individuals perceive a hollow mask as a normal face, schizophrenia patients do not as their low-level perception appears less constrained by top-down knowledge^24,25^. High precision of prediction error can explain perceptual organization deficits in schizophrenia, while high precision of prior beliefs^18,23^ has been postulated as a cause for hallucinations. Differences in precision weighting of priors and prediction errors across different cortical hierarchies may explain these contradictory accounts^19,26^: High-level association cortex has higher densities of recurrent connections^27^ favoring precision weighting of higher-order priors. The stage of the disease (early psychosis versus chronically medicated) may also play a role.

In synesthesia, exceedingly high precision for (top-down) priors may explain the persistent experience of concurrent sensations (e.g., color) despite the lack of actual sensory input (e.g., black letters)^28,29^. For synesthetes, we hypothesize an overreliance on prior beliefs^28^. Because synesthetes are aware their synesthesia is not real, it is suggested that it is mid-level priors that have high precision and reduce sensory prediction error, and not high-level priors. Another hypothesis is that synesthesia arises because prior predictions are too specific or detailed^30^.

### Approach

We evaluated the precision weighting of sensory signals and priors in perceptual inference in medicated chronic schizophrenia patients, synesthetes, and healthy controls. We used a well-established visual paradigm (Fig. 1A) which relies on perception of contours of letters or symbols in noise^31-33^. Letter recognition in literate individuals is guided by (implicit) long-term top-down priors^34^. In our task, the difference in noise density in and outside of the letter shape created vivid illusory contours, i.e. the subjective perceptual experience of contours in the absence of actual border lines^35^. Illusory contour perception, strongly influenced by previous experience, is believed to rely on cortical NMDA(N-methyl-D-aspartate)-dependent feedback projections^36,37^. The effect of top-down priors is strongest when sensory input is imprecise; i.e. weak, noisy or ambiguous^38^. We capitalized on this effect to investigate the precision weighting of sensory evidence and implicit long-term priors in perception across the three populations. We hypothesized that if precision weighting for sensory evidence is high compared to precision weighting for long-term priors, perception of illusory boundaries is less likely and visibility is low. This may occur in schizophrenia patients. In contrast, if long-term priors are dominant (high precision priors), perception of illusory boundaries is more likely and concomitantly visibility should be higher. This may occur in synesthetes when confronted with stimuli inducing precise long-term priors, e.g., graphemes eliciting synesthesia (Fig. S1).

**Figure 1.**
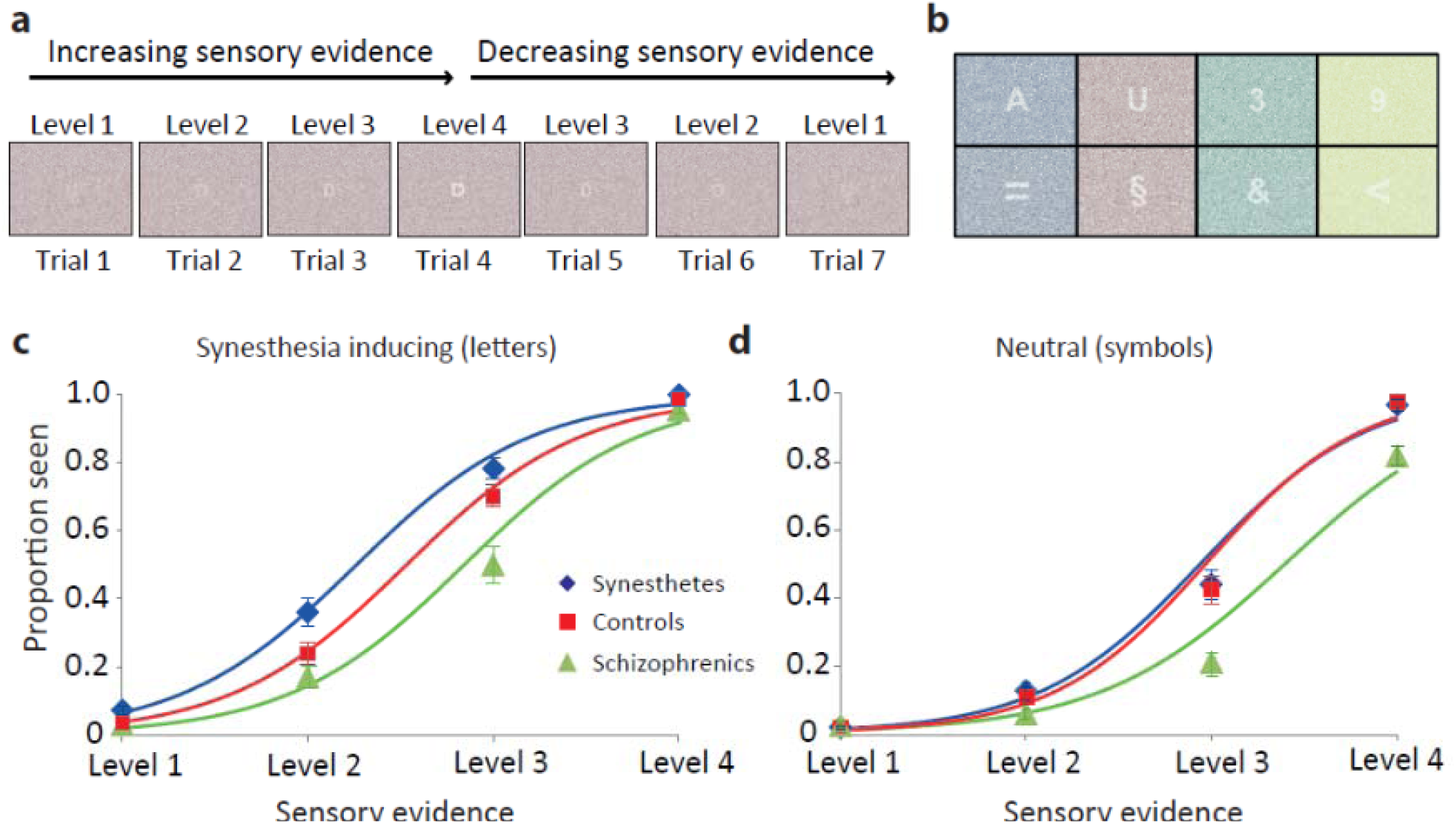
Paradigm and main results. (A) Example sequence, synesthesia-inducing stimulus condition. Across 7 trials, sensory evidence was first parametrically increased across 4 levels (trial 1-4) and then decreased (trial 5-7). The same token (letter, number, symbol) was used in a sequence. During sensory evidence increase, perception is strongly influenced by sensory evidence and implicit long-term top-down priors. In turn, when sensory evidence is decreased (trials 5-7), after stimulus recognition (around level 3-4), explicit, short-term top-down perceptual expectations further contribute to perception. (B) Example synesthesia (top) and neutral (bottom) stimuli. Stimulus background was either colored congruently with synesthetic stimuli, or randomly. (C and D) Psychometric curve fits for the synesthesia-inducing and neutral condition, respectively, for increasing sensory evidence trials (decreasing evidence phase results in Figure S2). Error bars depict the standard error of the mean. Results in main text.

To test these predictions, we parametrically manipulated sensory evidence (paradigm in Fig. 1A) while participants rated stimulus visibility. To leverage the relative precision of implicit long-term priors in synesthetes we presented either synesthesia-inducing stimuli (letters/numbers) or neutral stimuli (symbols) (Fig. 1B). Synesthesia-inducing graphemes were hypothesized to have long-term priors with higher precision weighting, which could in turn have a stronger effect on perception under conditions where sensory evidence is imprecise. These high-precision long-term priors are believed to convey additional color information about the stimulus (not ‘pop-out’), aiding letter identification because less sensory evidence is needed for inference to reach a sufficiently reliable solution. Thus, we expected long-term priors to selectively boost visibility for synesthesia-inducing stimuli in synesthetes, while the neutral condition served as an internal control for which no enhanced performance was expected. For schizophrenia patients, we hypothesized fewer perceived stimuli regardless of stimulus condition due to pervasive overreliance (higher precision weighting) on sensory evidence (Fig. S1).

Secondly, we evaluated whether additional short-term priors can normalize perception, i.e., bring schizophrenia patients’ and synesthetes’ perception closer to that of controls. For this purpose, we first increased sensory evidence (trials 1-4) until all stimuli were clearly recognized (Level 4 in Fig 1A); from this point on, an explicit, short-term, top-down prior was available. We then continuously decreased sensory evidence again to test this explicit prior’s effect (trials 5-7)^31-33^. This manipulation allowed us to separately investigate the contribution of *implicit*, long-term priors (trials 1-4) and *explicit*, short-term priors to perception (trials 5-7). While both affect perception their underlying mechanisms may differ.

## Methods

### Participants

Twenty synesthetes (mean age 29.2±8.2 years, 19 females), twenty chronic, medicated schizophrenia patients (mean age 39.7±12.4 years, 9 females), and twenty-six control participants (mean age 30.7±7.8 years, 17 females) participated in the study (demographics in Table 1). Study size was determined by calculating the required sample size for an expected medium-to-weak effect size at a power of 0.80 and α=.05 for a mixed-design repeated measures ANOVA with 3 groups^39^. A subset of 20 controls were specifically matched to the synesthete group in age and gender and an overlapping subset of controls was matched to the schizophrenia patients (all p>.10, see Table 1). Age and gender differed significantly between synesthetes and schizophrenia patients (*t*(38)=-3.08, p<.01; *t*(38)=-4.01, p<.001, respectively) and were included as covariates of no interest in all analyses. For additional recruitment, screening and synesthesia test details see Supplementary Methods. All participants gave written informed consent prior to the study, in accordance with the declaration of Helsinki. The study was approved by the ethics committee of the Medical Faculty at Goethe University Frankfurt, Germany.

**Table 1.**
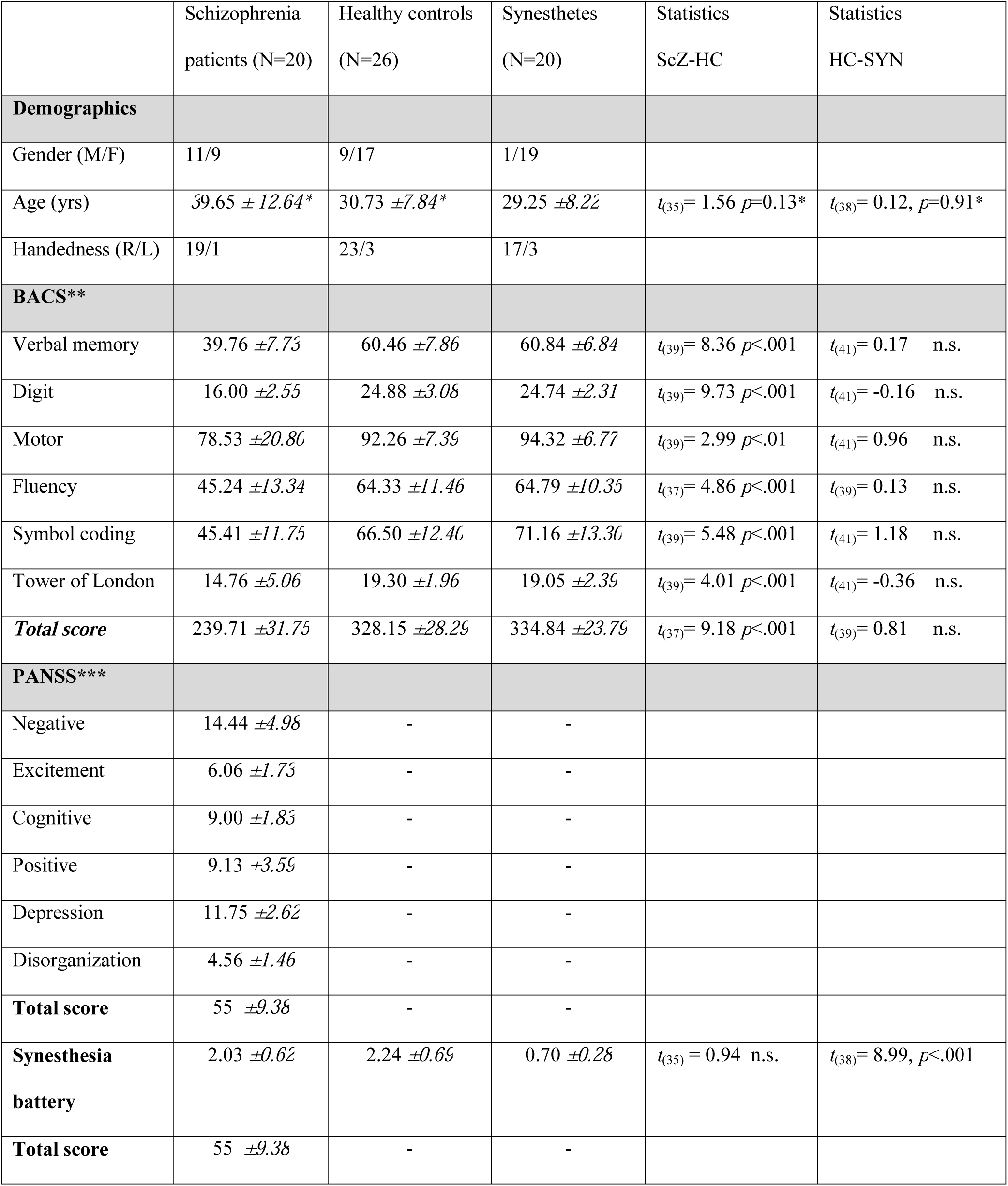
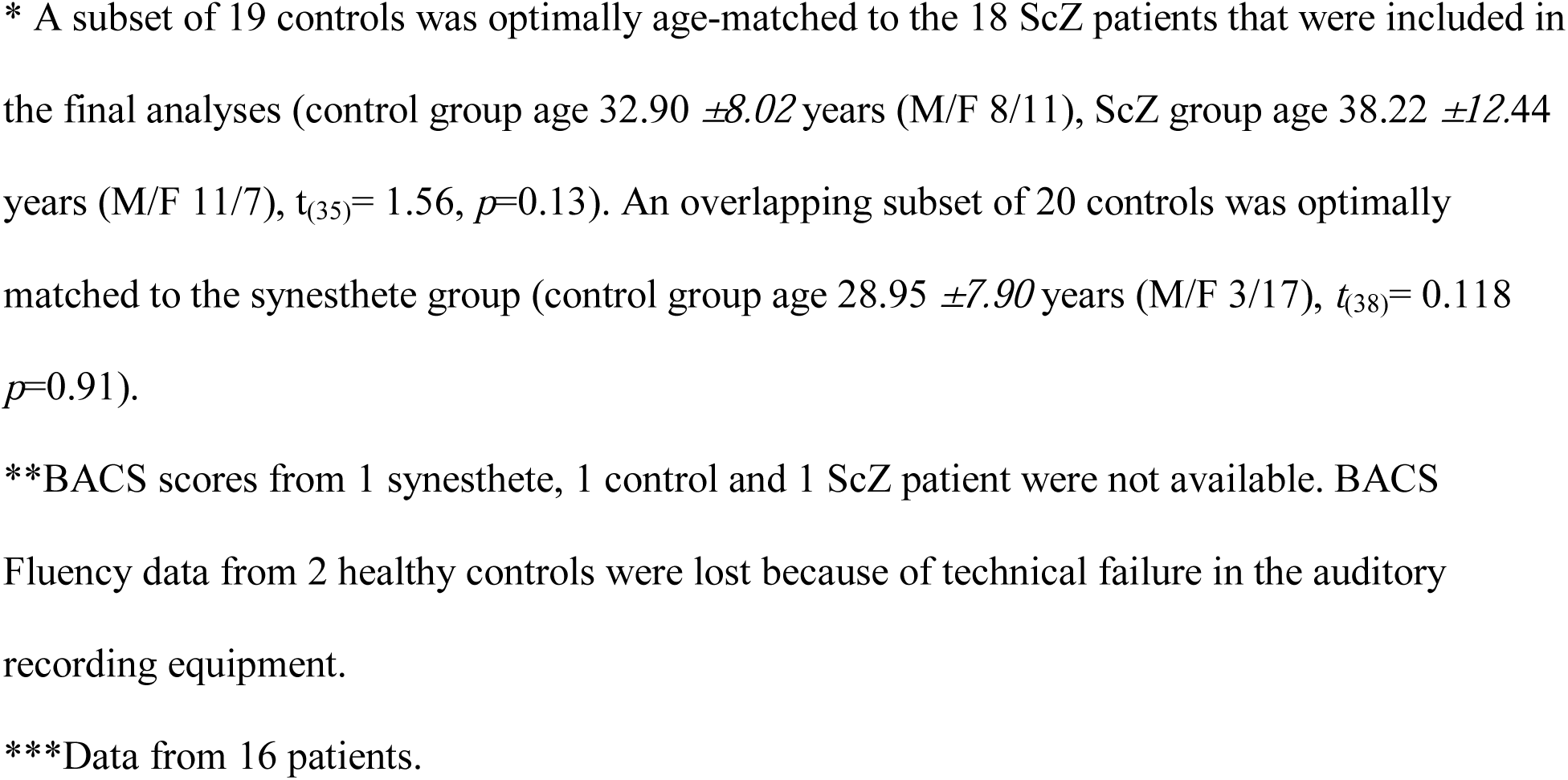
Demographics, PANSS and BACS scores of participants.

|  | Schizophrenia<br>patients (N=20) | Healthy controls<br>(N=26) | Synesthetes<br>(N=20) | Statistics<br>ScZ-HC | Statistics<br>HC-SYN |
| --- | --- | --- | --- | --- | --- |
| **Demographics** |  |  |  |  |  |
| Gender (M/F) | 11/9 | 9/17 | 1/19 |  |  |
| Age (yrs) | 39.65 ± 12.64* | 30.73 ± 7.84* | 29.25 ± 8.22 | $t_{(35)} = 1.56$ $p=0.13^*$ | $t_{(38)} = 0.12$, $p=0.91^*$ |
| Handedness (R/L) | 19/1 | 23/3 | 17/3 |  |  |
| **BACS\*\*** |  |  |  |  |  |
| Verbal memory | 39.76 ± 7.73 | 60.46 ± 7.86 | 60.84 ± 6.84 | $t_{(39)} = 8.36$ $p<.001$ | $t_{(41)} = 0.17$ n.s. |
| Digit | 16.00 ± 2.55 | 24.88 ± 3.08 | 24.74 ± 2.31 | $t_{(39)} = 9.73$ $p<.001$ | $t_{(41)} = -0.16$ n.s. |
| Motor | 78.53 ± 20.80 | 92.26 ± 7.39 | 94.32 ± 6.77 | $t_{(39)} = 2.99$ $p<.01$ | $t_{(41)} = 0.96$ n.s. |
| Fluency | 45.24 ± 13.34 | 64.33 ± 11.46 | 64.79 ± 10.35 | $t_{(37)} = 4.86$ $p<.001$ | $t_{(39)} = 0.13$ n.s. |
| Symbol coding | 45.41 ± 11.75 | 66.50 ± 12.40 | 71.16 ± 13.30 | $t_{(39)} = 5.48$ $p<.001$ | $t_{(41)} = 1.18$ n.s. |
| Tower of London | 14.76 ± 5.06 | 19.30 ± 1.96 | 19.05 ± 2.39 | $t_{(39)} = 4.01$ $p<.001$ | $t_{(41)} = -0.36$ n.s. |
| **Total score** | 239.71 ± 31.75 | 328.15 ± 28.29 | 334.84 ± 23.79 | $t_{(37)} = 9.18$ $p<.001$ | $t_{(39)} = 0.81$ n.s. |
| **PANSS\*\*\*** |  |  |  |  |  |
| Negative | 14.44 ± 4.98 | - | - |  |  |
| Excitement | 6.06 ± 1.73 | - | - |  |  |
| Cognitive | 9.00 ± 1.83 | - | - |  |  |
| Positive | 9.13 ± 3.59 | - | - |  |  |
| Depression | 11.75 ± 2.62 | - | - |  |  |
| Disorganization | 4.56 ± 1.46 | - | - |  |  |
| **Total score** | 55 ± 9.38 | - | - |  |  |
| **Synesthesia**<br>**battery** | 2.03 ± 0.62 | 2.24 ± 0.69 | 0.70 ± 0.28 | $t_{(35)} = 0.94$ n.s. | $t_{(38)} = 8.99$, $p<.001$ |
| **Total score** | 55 ± 9.38 | - | - |  |  |

\* A subset of 19 controls was optimally age-matched to the 18 ScZ patients that were included in the final analyses (control group age $32.90 \pm 8.02$ years (M/F 8/11), ScZ group age $38.22 \pm 12.44$ years (M/F 11/7), $t_{(35)} = 1.56$ , $p = 0.13$ ). An overlapping subset of 20 controls was optimally matched to the synesthete group (control group age $28.95 \pm 7.90$ years (M/F 3/17), $t_{(38)} = 0.118$ , $p = 0.91$ ).
\*\*BACS scores from 1 synesthete, 1 control and 1 ScZ patient were not available. BACS Fluency data from 2 healthy controls were lost because of technical failure in the auditory recording equipment.
\*\*\*Data from 16 patients.

### Stimuli and Design

We used a perceptual closure task^31-33^ in which participants viewed letters, numbers and symbols embedded in a colored noise background. Sensory evidence was parametrically manipulated by varying the noise level of the stimulus (Fig. 1A) with respect to its background in four steps, effectively providing contours for figure-ground segregation thereby controlling for stimulus’ visibility. Sensory evidence was first increased during the first 4 trials, increasing visibility, and subsequently decreased during trials 5-7, decreasing visibility. The same token (letter, number or symbol) was used across this 7-trial sequence. Perception is dominated by bottom-up input and long-term priors during the initial, *sensory evidence increasing phase* of the sequence (trials 1-4). In the subsequent sequence, the *sensory evidence decreasing phase*, when subjects have recognized the stimuli, short-term top-down priors aid recognition^31-33^.

Stimuli used in the sequences were such that in synesthetes they could either elicit synesthesia (letters and/or numbers), or be neutral, not eliciting synesthesia (symbols). For controls and schizophrenia patients none of the stimuli elicited synesthesia. Thus, this manipulation was – although identical in all three groups – only relevant for synesthetes and used to manipulate their long-term priors. All stimuli were embedded in a colored noise background. In the synesthesia inducing condition, the background color was congruent with the synesthetic color of the stimuli for that particular synesthete. One or more controls and one schizophrenia patient also viewed that same physical stimulus list (see Supplementary Material). The background color was also randomly assigned to tokens of the neutral, non-synesthetic condition, as symbols did not elicit synesthesia. This prevented precuing a condition (i.e., synesthetic or non-synesthetic condition). Example stimuli for both conditions are shown in Fig. 1B. Further details about stimulus selection and presentation are provided in the Supplementary Material.

A total of 1260 trials were presented in four experimental blocks. On each trial (Fig. 1A and Supplementary Fig. 2A), participants rated the subjective visibility of the stimuli on a 4-point Perceptual Awareness Scale (PAS)^40^ defined as 1. No experience; 2. Brief glimpse; 3. Almost clear impression; 4. Clear impression. The use of the PAS was rehearsed with the participants in a practice session of 5-10 sequences to ensure that participants grasped the difference between level 2 (not visible) and 3 (visible) responses. To evaluate the use of the PAS by the participants and false alarm rate, clearly visible (Level 4) and clearly invisible (Level 1) stimuli where added to the sequence.

**Figure 2.**
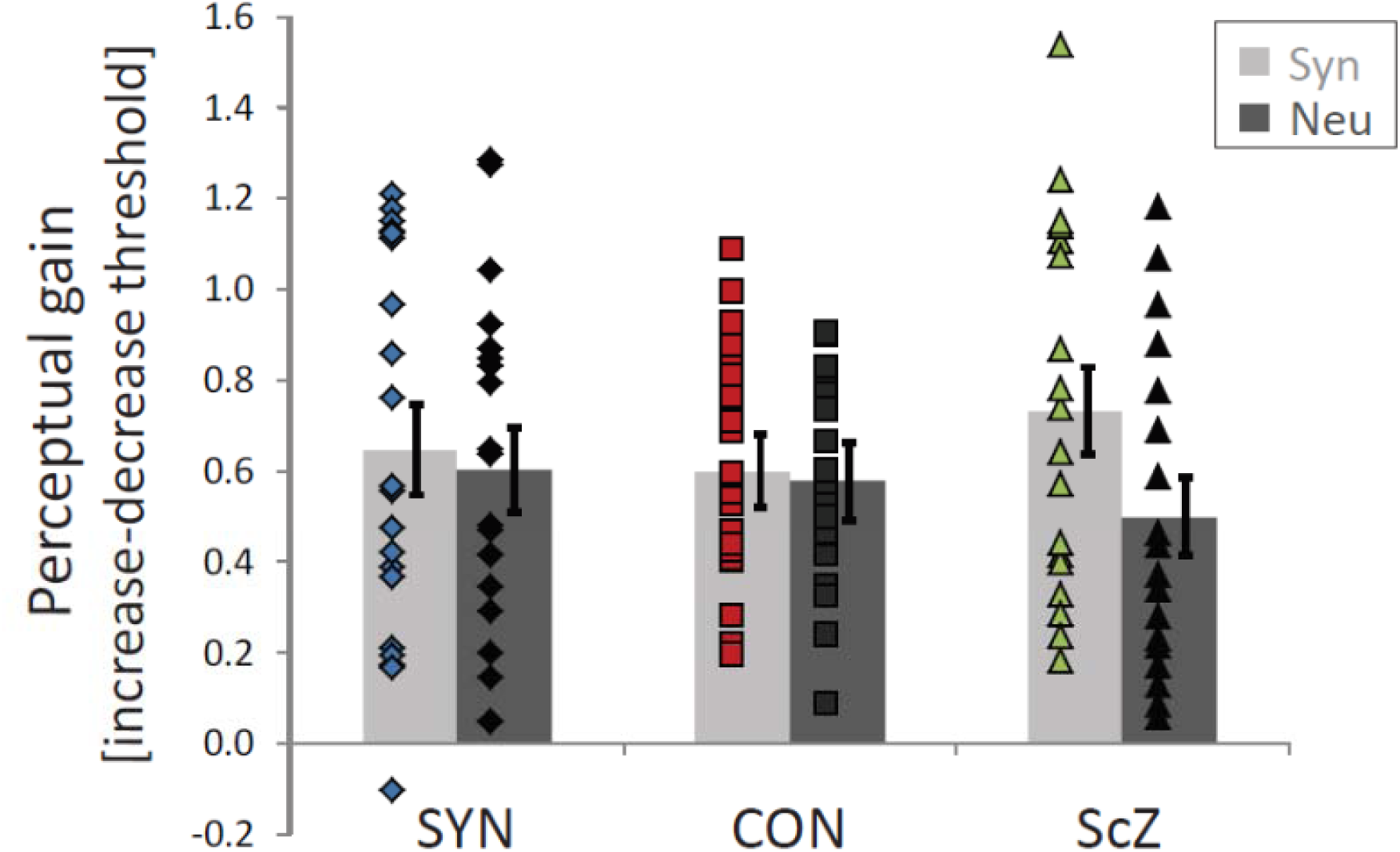
Explicit priors boost perception. Stimulus recognition creates an explicit short-term prior, lowering perceptual thresholds (trials 5-7) even though sensory evidence remains equal (to trials 1-3). Perceptual gain is plotted (increasing minus decreasing phase perceptual threshold) and is similar across groups. Diamonds, squares, and triangles represent single participants. Error bars depict the standard error of the mean. SYN, synesthetes; CON, controls; ScZ, schizophrenia patients, Syn, synesthesia condition; Neu, neutral condition.

### Fitting of psychometric functions

For analyses, responses were recoded into a visibility measure of recognition with categories: Not visible (responses 1 and 2) or Visible (responses 3 and 4). The resulting psychometric data from each participant and condition was fitted with logistic functions, each defined by three parameters: threshold, slope, and fixed lapse-rate^41^, using the Palamedes Toolbox (Version 1.6.0) for Matlab (http://www.palamedestoolbox.org/). Guess rates were fixed at 0.5 (i.e., chance level) across all subjects and conditions. All three parameters were fitted separately per subject and stimulus condition. Quality of fit for each subject was determined by assessing the square sum of the errors.

### Statistical analysis

Fitted threshold values (inferred stimulus level at 50% seen stimuli) for each participant were submitted to a mixed-repeated measures ANOVA with Group as a between-subject factor (synesthetes, controls, schizophrenia patients) and as within-subject factors Stimulus Condition (synesthesia inducing/neutral) and Phase (sensory evidence increase/sensory evidence decrease). Age and gender were included as covariates of no interest.

Data from 4 subjects were removed prior to statistical analysis because they performed 2 SD below their group mean (2 controls, and 2 schizophrenia patients). For 1 control, curve fitting failed to provide a valid threshold value in the decreasing sensory evidence conditions because the participant continued to rate the stimuli as highly visible and never reached the 50% visibility threshold; this control was also removed from analyses. Five schizophrenia patients completed less than 4 blocks of the experiment due to fatigue. All reported tests are 2-sided with α=0.05.

## Results

Comparing perceptual thresholds across groups, we observed a group effect (*F*(2,56)=6.89, *p*=.002, η _p_^2^=.20) modulated by whether stimuli induced synesthesia (Group x Stimulus condition *F*(2,56)=6.76, *p*=.002, η _p_^2^=.19, Fig. 1C-D). Compared to controls, schizophrenia patients perceived fewer stimuli during the increasing sensory evidence phase (group effect *F*(1,37)=11.90, *p*=.001, η _p_^2^=.24) both for synesthesia-inducing (*F*(1,37)=6.63, *p*=.014, η _p_^2^=.15) and neutral stimuli (*F*(1,37)=16.83, *p*<.001, η _p_ ^2^=.31), indicating more ‘veridical perception’ and suggesting an overreliance on sensory evidence. Synesthetes selectively perceived more stimuli than controls in the synesthesia-inducing condition during sensory evidence increase; there was no group effect for the neutral condition (Group x Stimulus condition *F*(1,39)=9.12, *p*=.004, η_p_^2^ =.19; Group effect synesthesia-inducing *F*(1,39)=10.62, *p*=.002, η_p_^2^ =.21; Group effect neutral *F*(1,39)=1.59, *p*=.215). This suggests that synesthetes profited specifically from an additional implicit, long-term prior (synesthetic color). The spatial experience of synesthesia i.e., projector/associator subtype, did not modulate thresholds (see Supplementary Results). These results are not explained by differences in response criterion across the populations as indicated by a low and comparable number of false alarms for stimuli that do not support perception (level 1; see Supplementary Results).

We also evaluated the contribution of *explicit, short-term* priors to perception. We found they boosted perception (*F*(1,56)=5.77, *p*=.020, η_p_ ^2^=.093) by lowering perceptual thresholds in the sensory evidence decreasing condition similarly across all groups (*F*(2,56)<1, n.s., Fig. 2). However, they did not override the different perceptual phenotypes from the increasing sensory evidence phase (Supplementary Fig. 2B/C; non-significant Stimulus condition x Phase x Group interaction (*F*(2,56)=2.86, *p*=.066, η_p_^2^=.093).

The selective reduction in visibility threshold for the synesthesia-inducing condition in synesthetes is consistent with the hypothesis of stronger, high precision implicit long-term priors. However, strategic usage of the colored background by the synesthetes could yield comparable results. To rule this out, we investigated learning of the stimulus set; and specifically a differential learning effect by the synesthetes as compared to the other two groups. We analyzed the visibility scores and reaction times (RTs) for the first three instances of each stimulus for synesthetes (*N*=20) and their matched controls (*N*=20) (for details see Supplementary Methods). While learning is common during experiments and a main effect of repetition was anticipated, an interaction between group, condition, and repetition would indicate learning occurs at a differential rate for synesthetes (specifically for the synesthesia-inducing condition, i.e., letters/numbers).

For visibility scores, none of the interactions with Repetition were significant (all *p*>.10). As expected, we found a main effect of Repetition (*F*(2,76)=15.9, *p*<.001, η_p_^2^=.30), indicating stimuli were better recognized with repeated occurrences. A Group x Condition interaction was also present (*F*(1,38)=7.32, *p*=.010, η_p_^2^=.16) confirming our main results of better performance for the synesthetes specifically in the synesthesia-inducing condition. For RTs, we only observed a Repetition x Group interaction, *F*(2,76)=4.00, *p*=.022, η _p_^2^=.095), whereby synesthetes exhibited a repetition effect (*F*(2,38)=9.20, *p*=.004, η_p_ ^2^=.33) while controls did not (*F*(2,38)=1.39, *p*=.26, η_p_^2^ =.068). Critically, the effect of repetition on the RTs in synesthetes was not modulated by whether the stimuli elicited a synesthetic experience (interaction of Repetition x Condition (*F*(2,38)<1, n.s.)) and there was also no Group effect in the RTs (*F*(1,38)=0.67, *p*=.42). Thus, synesthetes do not appear to make explicit, strategic use of their synesthetic color which could explain their lowered psychophysical thresholds. Additional support comes from the analysis of RTs for synesthesia-inducing stimuli upon the first stimulus encounter, which shows that synesthetes are not faster than controls to recognize synesthesia-inducing stimuli at first presentation (see Supplementary Results).

We explored whether performance was influenced by synesthetic consistency, by positive/ negative symptoms (PANSS scores) and/or cognitive performance (BACS scores). These correlations are exploratory given the limited sample size. For synesthetes, consistency scores did not correlate significantly with performance in any condition (all *r*(19)<.320, all *p*>.18). For schizophrenia patients, the Cognitive PANSS subscore correlated with performance on the synesthesia-inducing sensory evidence decreasing condition (*r*(16)=.498, *p*=.050, 95% CI [.293,.733]), and marginally with the synesthesia-inducing sensory evidence increasing condition (*r*(16)=.483, *p*=.058, 95% CI [.117,.769]), but these effects did not survive correction for multiple comparisons. BACS performance did not correlate significantly with visibility thresholds in schizophrenia patients. The strongest correlation was observed between the Total BACS score and the visibility threshold in the neutral sensory evidence increasing condition, but was not significant (*r*(17)=-.418, *p*=.095, 95% CI [-.732, .133]).

## Discussion

The results on perceptual thresholds support the hypothesis that imbalances in perceptual inference may underlie different perceptual phenotypes. While the performance of schizophrenia patients on our perceptual closure task can be explained by an overreliance on sensory evidence, i.e. high precision prediction errors, synesthetes appeared to profit from high-precision long-term priors. Analysis on the learnability of the stimulus set rules out the possibility that the lowered visibility threshold in synesthetes may be explained by strategic control reinforcing the proposal that enhanced precision of long-term priors specific to that population may explain the differences in perceptual threshold.

How are differences in perceptual inference manifested in neural mechanisms? For schizophrenia, dysfunction of NMDA-receptors has been implicated in disease progression and symptoms^42^. Glutamatergic NMDA-receptor mediated signaling is involved in top-down predictive signals, inhibiting incoming sensory information^19^. Hence, dysfunction of NMDA-receptors potentially explains our results by weakening predictive signals, leading to aberrant high precision weighting of sensory prediction errors. Additionally, dopaminergic signaling is implicated as a neuromodulator of precision weighting^43^, enhancing the precision (saliency) of bottom-up signals on the basis of reward. When considering the underlying mechanisms we note that our patients were medicated with anti-dopaminergic medication, and that NMDA-receptor functioning was not measured explicitly.

Synesthesia has traditionally been explained by cross-activation and disinhibited feedback accounts^8^, explaining brain activity for concurrent synesthetic experiences through anatomical cross-wiring or disinhibited feedback signals, respectively. In terms of predictive processing, Seth^28^ has emphasized the role of high-precision mid-level priors in maintaining synesthetic experience in the absence of sensory input. An alternative predictive processing account hypothesizes synesthesia arises when (categorical) predictions are too detailed, increasing bottom-up prediction error because a lot of variance goes unexplained by prior predictions^30^. Our results are in concordance with high precision priors in synesthetic experience. Given that synesthetes performed as controls on the neutral trials, our results provide no evidence of generally altered perceptual inference in synesthetes. We propose the high-precision priors for synesthesia inducing stimuli originate beyond early visual cortex (Fig. S1).

Remarkably, while clearly differing in the weighting of long-term priors, schizophrenia patients and synesthetes profit equally from additional, short-term top-down knowledge to improve perception. Yet, those explicit, short-term cues do not normalize perception neither in schizophrenia patients nor in synesthetes relative to controls. This suggests that additional short term priors derived from context are not strong enough to overcome the fundamental imbalance in precision weighting that is present in perceptual inference in both groups. Short term priors may be task-related, relatively low-level, allowing for performance-optimizing for both participant groups. Simultaneously, the overall imbalance in precision weighting of sensory evidence versus high-level priors is not altered during the experiment. Dopaminergic signaling is implied in determining the level of precision and may be affected in schizophrenia^44^.

It should be noted that the effects of priors on perception may depend on the specific task that was performed and on the processing hierarchy of the stimuli that were involved^45^. In the current study the long-term priors of letters and symbols are Gestalt-like priors; yet the task is a low-level task involving contour detection. For schizophrenia, low-level processing has been associated with a decreased influence of priors (e.g.^19^), explaining insensitivity to illusions that are driven by long-term priors such as illumination priors^19,24,25^. On the contrary, for hallucinations, which are positive symptoms associated with schizophrenia, a stronger reliance on prior information has been reported^18^. As Sterzer^19^ proposes, weak priors at low-level may lead to perceptual uncertainties that may be compensated by reliance on high-level abstract or semantic prior beliefs, which in turn lead to hallucinations. Similarly, indeed for synesthetes an increased influence of priors was only seen in the specific task conditions where synesthesia was present. Thus, when generalizing the effects of priors on perception to other studies, processing hierarchy and task demands should be taken into account^19,45^.

Aberrant precision weighting is hypothesized to underlie psychopathology in the wider sense^20^, e.g. also including autistic perception^46-48^. In autism, high-precision, inflexible prediction errors are proposed, while it is debated whether priors are overfitted or too weak^47,48^. This explains reduced top-down influences in perception in autism^49^. Synesthesia is more prevalent in autism^50,51^, with similar perceptual profiles^11,52^, and overfitting of (high-precision) priors is also hypothesized for synesthesia^28^. Synesthesia is associated with positive, but not negative, schizotypy^53,54^. We focused on perception, not on positive symptoms, and our results clearly speak in favor of different perceptual phenotypes in synesthetes and schizophrenia patients, despite any potential overlap.

Participants reported subjective stimulus visibility without performing an objective discrimination task. Therefore, it is theoretically possible that the visibility ratings used for curve fitting did not reflect objective stimulus visibility. In fact, we have previously demonstrated that visibility rating and objective performance can dissociate^55^. Thus, our study concerns differences in subjective perceptual phenotypes, not addressing alterations in objective performance. Our task captures subjective perceptual abilities and does not merely reflect response bias^32^, see also Supplementary Results.

Another study limitation is the schizophrenia sample size (N=18). By using a sensitive experimental design, we were able to detect differences with a medium-to-weak effect size. Our schizophrenia patients were medicated and they had an average disease duration of 14 years, qualifying them as chronic patients, representative of the majority of schizophrenia patients. The anti-dopaminergic medication, however, may influence precision weighting and retinal functioning^44,56^. A strength of our study is the inclusion of three clearly distinct groups allowing us to draw conclusions across a variety of phenotypes. Schizophrenia patients as well as synesthetes were carefully characterized with regard to their specific conditions to ensure samples were as homogeneous as possible and not confounded by other factors (e.g. substance abuse, other diagnoses).

In summary, we demonstrate that for a variety of perceptual phenotypes – schizophrenia patients, controls, synesthetes – perceptual variability can be explained by differences in precision weighting of bottom-up sensory evidence and top-down priors. This clear behavioral profile might serve as a phenotypic characterization to be used in computational modeling and neurophysiological investigations in order to better characterize the neural mechanisms subserving these perceptual differences. Ultimately, this can contribute to a better understanding of the pathophysiology of schizophrenia.

## Acknowledgments

We thank Caspar M. Schwiedrzik for invaluable comments on the manuscript and Dr. Michael Grube for assistance in schizophrenia patient recruitment.

## Author Contributions

T.M.v.L. performed experiments; L.M. conceptualized the study; T.M.v.L. and L.M. designed the experiments, analyzed the data, and wrote the paper; A.S. assisted in patient recruitment and behavioral testing; A.-M.J. assisted in participant recruitment; M.W. provided data acquisition resources; P.J.U. assisted in patient recruitment and institutional review board approval; W.S. reviewed and edited the paper. W.S. and L.M. secured funding.

