## Supplementary Material for "Perceptual phenotypes: Perceptual gains and losses in synesthesia and schizophrenia"

*Contents:*

**Supplementary Figure 1 and 2**

**Supplementary Material and Methods**

**Supplementary Results**

**Supplementary References**

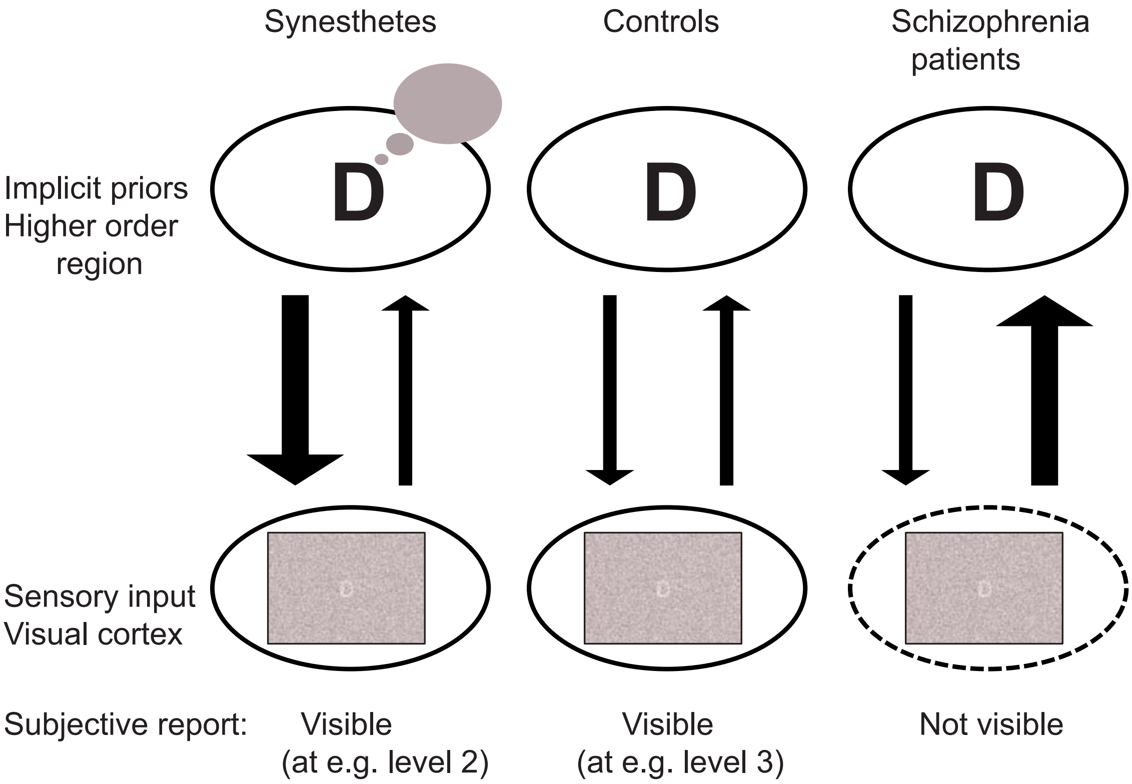


**Figure S1 Hypotheses for the different groups**

We hypothesize that implicit prior knowledge (in higher order brain regions) and sensory evidence (bottom up signals) are weighted differently across the three participant groups, leading to differential perceptual phenotypes (subjective reports) even though the stimuli that are presented to all participants are the same. Depicted is the condition for the trial sequence providing increasing sensory evidence at a stage where explicit priors are not yet available, and synesthesia inducing stimuli are used (a letter). In evaluating the visibility of the stimuli, synesthetes may be aided by precise, implicit long-term priors (thick downward arrow) that also hold a representation of the color of the letter (D = dark pink). In schizophrenia patients, precision of sensory evidence may be higher (thick upward arrow) while precision of implicit long term priors is reduced (thin downward arrow). For controls, precision of sensory evidence and implicit long-term priors are equal.

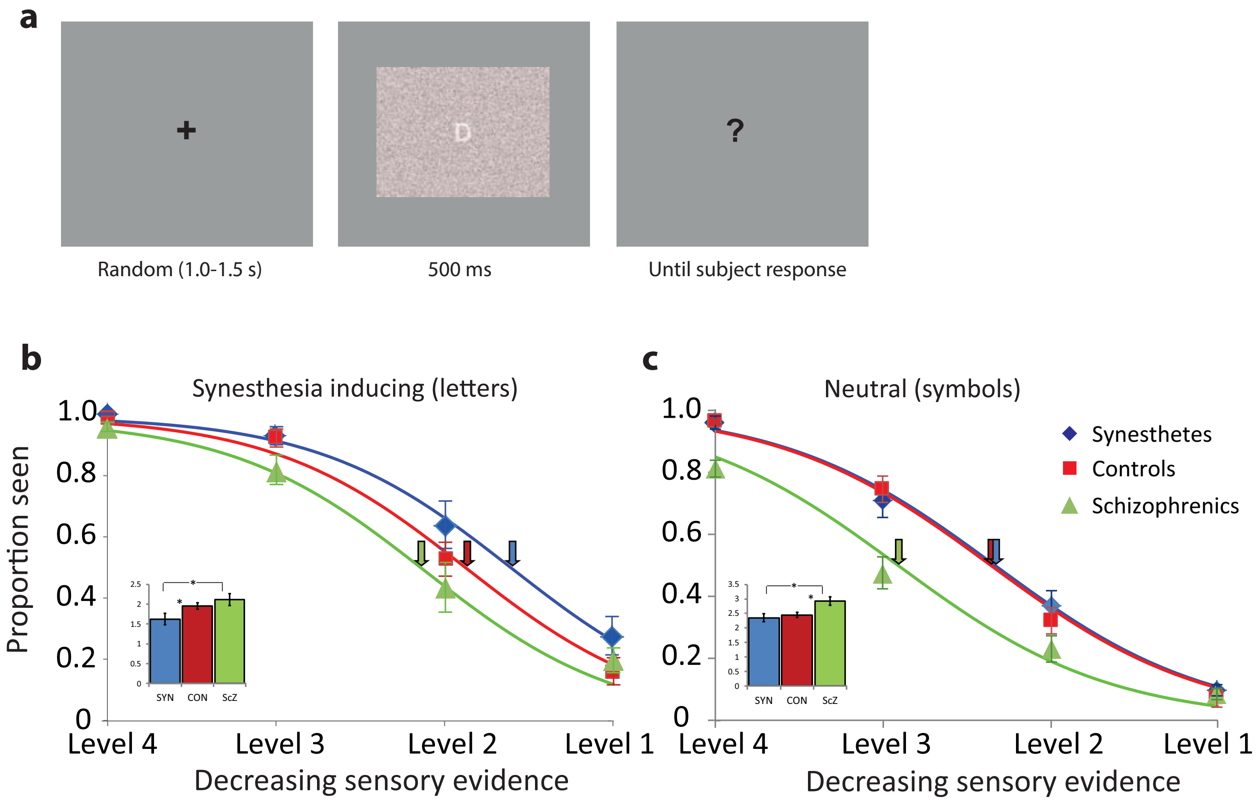


**Supplementary Figure 2. Trial sequence and decreasing sensory evidence results**

(A) Time course of one trial within a sequence. A trial began with an inter-trial interval of random duration (1-1.5 secs), during which a fixation cross was presented. Next, the stimulus was displayed for 0.5 secs, followed by a question mark prompting participants to indicate the visibility of the stimuli according to the PA Scale[1]. (B and C) Psychometric functions for all three groups during decreasing sensory evidence trials, for synesthesia-inducing (B) and neutral (C) stimulus conditions. Explicit priors aid perception in all groups as evidenced by a lower threshold of visibility in the decreasing sensory evidence condition. However, thresholds for perception (see insets in B and C) are nonetheless higher in schizophrenia patients compared to the other groups, especially for the neutral condition (synesthetes vs schizophrenics: 2.35 vs 2.93, $t(36)= -3.00$, $p<.01$; controls vs schizophrenics: 2.45 vs 2.93, $t(39)= -3.13$, $p<.01$). In synesthetes, threshold for synesthesia inducing stimuli remains lower than for the other two groups (synesthetes vs schizophrenics: 1.62 vs 2.11, $t(36)= -2.40$, $p<.05$; synesthetes vs controls: 1.62 vs 1.95, $t(30.12)= -2.13$, $p<.05$ (df adjusted for inequality of variance). Pointing arrows correspond to the perceptual threshold, colored differently for each group. Error bars depict standard error of the mean.

### Supplementary Material and Methods

### Participants

All participants completed a short screening questionnaire about their medical history and a test for synesthesia prior to participation (the "Synesthesia Battery"[2]; explained below, for scores see Table 1). Synesthetes additionally completed two questionnaires about the spatial location of their synesthesia: the Projector-Associator questionnaire[3,4], and the Illustrated Synesthetic Experience Questionnaire[5]. Controls and synesthetes reported no history of neurological or psychiatric disease, no medication use at the time of the study, and normal or corrected-to-normal vision. Chronic schizophrenia patients were recruited from the psychiatric out-patient unit of the Clinic Frankfurt-Höchst. All patients fulfilled DSM-IV criteria for schizophrenia as verified by a trained psychologist by means of a Structural Clinical Interview for DSM-IV-R (SCID) prior to inclusion. Average disease duration was 14.1±12.9 years; all patients were medicated with atypical neuroleptics. Current psychopathological symptoms were assessed using the Positive and Negative Syndrome Scale (PANSS) for schizophrenia[6,7]. Scores were obtained for Negative, Positive, Excitement, Cognitive, Depression, and Disorganization subscales (see Table 1). Cognitive function was assessed with the German version of the Brief Assessment of Cognition in Schizophrenia (BACS) in all subjects[8]; for scores see Table 1.

*Synesthete recruitment and inclusion*

Synesthetes were recruited by advertisement of the study via email to students in Frankfurt am Main and surrounding cities, by advertisement on the internet, and from earlier studies. Prior to inclusion in the study, developmental grapheme-color synesthesia was established by means of a synesthesia questionnaire (e.g. ‘‘How long have you experienced synesthesia?’’)[4] and by online completion of the standardized grapheme-color subtest of the Synesthesia Battery, which evaluates the consistency of synesthetic experiences[2]. Consistency is a defining characteristic of the condition[9] (but see also[10,11]). In the Synesthesia Battery, subjects are presented with all 26 letters of the alphabet and digits 0-9 three times in random order, and requested to choose from a color palette, the color (on a red, green, blue [RGB] scale) that best matches their specific synesthetic experience. Differences in RGB value between the three instances of each grapheme are used to compute a difference score. Difference score values below 1.0 are considered to signal consistent synesthetic color experiences in which variation between the three instances of the color is low[2] (but see[12] for argumentation that a more lenient cut-off score of 1.43 also suffices). All twenty synesthetes except one scored below 1.0, the average score was 0.70±0.28. One synesthete reported multiple simultaneous colors for several graphemes, therefore exhibiting less consistency during the Battery test (score of 1.48), but was still included in the study on the basis of her subjective reports in the synesthesia questionnaire. One synesthete did not complete the Synesthesia Battery but completed a regular grapheme-colour 36 item test-re test synesthesia consistency task on which she scored 31 items correct (88.6%).

*Control recruitment and inclusion*

Neurotypical participants were recruited locally in Frankfurt am Main via online advertisement and word of mouth. In addition to the standard (medical) screening, controls were interviewed about synesthetic experiences and completed the grapheme-color test of the Synesthesia Battery online to exclude synesthesia. Controls were well above the 1.0 threshold for synesthesia and scored, on average, 2.24±0.69 on the Synesthesia Battery. Three controls did not complete the Battery and one control chose only black for all graphemes, not yielding a meaningful consistency score. No difference in the synesthesia battery scores were found between controls and schizophrenia patients ($t(35) = 0.937$, $p=.36$).

*Schizophrenia patient synesthesia characteristics*

Schizophrenia patients were additionally interviewed about possible synesthetic experiences and completed the grapheme-color test of the Synesthesia Battery at the laboratory to exclude synesthesia. In this case the Synesthesia Battery was run in the offline Matlab version of the toolbox. On average, patients scored 2.03±0.62 on the Synesthesia Battery. One patient scored within the synesthetic range for numbers and vowels only (score of 0.56) and one patient completed a spoken test-re-test letter-colour consistency task instead of the Battery (score of only 4% consistent). Because priority was given to completion of the main experiment, 4 patients did not complete a synesthesia test.

### Stimulus selection and presentation

Ten letter/digit stimuli eliciting strong synesthetic experiences, with maximum synesthetic color variety, and sufficient variety in visual shape were selected separately for each synesthete on the basis of their synesthetic color experiences and used for the synesthesia condition. Additionally, ten non-synesthesia inducing symbols were chosen for each synesthete and used for the neutral condition. Prior to the experiment, synesthetes selected their individual synesthetic colors for the ten letters/digits on the experimental computer to ensure proper matching of the stimulus colors to the actual synesthetic color of the stimuli. Stimuli were then generated separately for each synesthete using custom code in Matlab (Mathworks, http://mathworks.com).

A total of 1260 trials were presented in 180 sequences of 7 trials. Ninety sequences belonged to the synesthetic-inducing condition (9 sequences per letter) and ninety sequences to the neutral condition (9 sequences per symbol). Subjects performed four blocks of 45 sequences each, with self-paced breaks between blocks. The experiment took approximately 80 minutes to complete. Each trial (Supplementary Fig. 1) started with a random inter-trial interval (1.0-1.5 secs) during which a black fixation cross on a grey background was displayed. The stimulus was then shown for 0.5 secs immediately followed by a question mark presented until the subject's response. The subject response prompted the next trial.

Stimuli were presented in pseudo-randomized order using Presentation (Neurobehavioral Systems, https://www.neurobs.com/). Two consecutive sequences did not display stimuli either of the same color, or the same token. A maximum of 5 consecutive sequences from the same condition was permitted (synesthetic inducing or neutral), and two different font sizes per stimuli were used to avoid adaptation and/or perceptual learning. Stimuli of the same font size repeated maximally 5 times in a row.

Controls ($N$=26) and schizophrenia patients ($N$=20) received one of the 20 unique stimulus lists previously generated for each of the 20 synesthetes (conserving the same physical stimuli and stimulus order). This was done since unique stimuli had to be used for each synesthete, introducing variability in stimulus shape for each participant in addition to varying degrees of discriminability due to the different background colors (e.g. visibility against a dark blue or bright yellow background, see also Fig. 1B). Matching the stimulus lists between the groups removed these confounds from our dataset. Since there were more controls than synesthetes, several stimulus lists were used twice for the control group.

Magnetoencephalography (MEG) data were collected during the study; hence participants performed the experiment while being seated in an electrically shielded and sound attenuated room. Stimuli were presented on a transparent screen with a grey background located 51 cm in front of the subjects. An LCD projector (60 Hz refresh rate) located outside the magnetically shielded room was used to project the stimuli onto a screen inside the MEG cabin via 2 front-silvered mirrors. The grey background (Figure S1) measured 29.1 x 37.9 degrees of visual angle and the stimulus display measured 22.6 x 30.2 degrees of visual angle. Grapheme stimuli themselves were created using the same font and measured maximally 3.9 x 3.6 degrees of visual angle depending on the exact letter/digit/symbol shape.

**Learning of the stimulus set**

To evaluate possible learning across the stimulus set during the experiment we analyzed visibility scores and reaction times (RTs) for the first three instances of each stimulus separately for synesthetes ($N$=20) and their matched controls ($N$=20). RTs below or above 2.5 SD from the mean for each participant and condition were removed prior to analysis. Visibility scores and RTs were subjected to a mixed repeated measures ANOVA with the between-subject factor Group (synesthetes/controls) and the within-subject factors Stimulus condition (synesthesia inducing/neutral), Phase (sensory evidence increase/sensory evidence decrease), Repetition (first, second, and third occurrence of the stimuli) and Stimulus level (1,2,3). For results see main text.

### Supplementary Results

*Psychophysical thresholds are not modulated by the spatial location of synesthetic experience.*

Synesthetes differ in the spatial location where they experience synesthesia, and have been classified as projectors or associators based on these differences. 'Projector' synesthetes tend to experience their synesthetic colors in the outside world, often located at the place where the inducer is located. 'Associator' synesthetes experience their synesthesia as strong associations, e.g. 'in the mind's eye'. Projector and associator synesthetes differ in various behavioral and physiological measures and projector-associator status may influence experimental outcomes[13-15]. We therefore verified that our main findings were not restricted to only projector or only associator synesthetes.

We assessed the projector-associator status of our population of synesthetes by means of two questionnaires: the Projector-Associator questionnaire (PAQ)[3,4], consisting of ten statements related to the synesthetic experience for which participants indicated to which extent they agreed to the statement on a 5-point Likert scale; and the Illustrated Synesthetic Experience Questionnaire (ISEQ) which makes use of visual illustrations of what the synesthetic experience could be like[5]. Based on the responses to these questionnaires, which correlated strongly ($r$=0.873, $p$<.001, 95% CI [.761, .944]), we classified 8 subjects as projectors and 11 as associators; for one person the status could not be determined with certainty. We correlated the PAQ-score with the consistency of the synesthetic experiences, and with the perceptual thresholds in all four experimental conditions (synesthesia inducing sensory evidence increasing, synesthesia inducing sensory evidence decreasing, neutral sensory evidence increasing, neutral sensory evidence decreasing) to investigate whether projector-associator status affected any of the experimental outcomes. No significant effects were found (all $p$>.37), indicating that the location of the experienced synesthetic colors did not influence task performance.

*Synaesthetes do not make strategic use of the color to detect stimuli*

To further evaluate whether synesthetes make explicit use of the color priors to aid grapheme recognition, we also evaluated whether synesthetes recognize synesthesia inducing stimuli faster than neutral stimuli upon the first encounter. We evaluated differences in RT between conditions for the first encounter of the stimuli for synesthetes ($N$=20) in repeated measures ANOVA with the within-subject factors Stimulus condition (synesthesia inducing/neutral), and Stimulus level (1, 2, 3). Only data from the increasing sensory evidence phase were included.

When collapsing across all stimulus levels (1-3, leading to different degrees of visibility), we found comparable RTs for synesthesia inducing and for neutral stimuli (levels 1-3, $F(1,19)=2.92$, $p=.104$, $\eta_p^2=.13$). However, when we investigated the effect separately per stimulus level, we observed that for stimulus level 2, at the visibility threshold, RTs for synesthesia inducing stimuli were slower than for the neutral condition, 1201 ms and 997 ms respectively ($F(1,19)=7.46$, $p=.013$, $\eta_p^2=.28$). While for level 1 and 3, for which participants either do not perceive the stimuli due to the strong noise level or perceive them clearly due to low noise level, there was no difference in the RT ($p=.42$ and $p=.49$). Altogether, these results indicate that synesthetes do not make strategic use of the color cue to anticipate synesthetic inducing stimuli upon the first encounter of a stimulus. If anything, synesthesia inducing stimuli are recognized later than neutral stimuli. This RT difference for stimulus level 2 concurs with previous electrophysiological findings using the same paradigm reporting that synesthetes exhibited a delayed visual P200 peak response, exclusively for synesthesia inducing stimuli[16]. The P200 is an ERP component related to conscious visual processing[17,18]. The slowing of the EEG P200 component and the RTs in our experiment at level 2 for synesthesia inducing stimuli only may then reflect the extra processing time needed to bind the inducer to the synesthetic color, a process which is not needed for neutral stimuli which do not elicit synesthesia. As synesthetes recognized almost 40% of synesthesia inducing stimuli at threshold (level 2), this extra binding process could explain the slower RTs in our study.

*Response bias does not differ across populations*

The task used in this study has been carefully calibrated to rule out factors such as response bias. In particular, in two previous studies we demonstrated that memory-based predictions and not the mere sequential presentation of information explains the reduction in the visibility threshold; and also that visibility thresholds are reduced only in the presence of memory-based predictions that match sensory evidence, consistent with a bayesian optimal integration as opposed to a sticky response bias that inflates visibility artificially[18,19]. Moreover, the introduction of sensory levels that should not lead to clearly visible stimuli – level 1 – enables to directly investigate response bias, and in particular false alarms. To evaluate response biases across the three populations, we focused on level 1 and investigated the amount of false alarms i.e., proportion of trials in which subjects declared to have 'seen' (PAS scales 3 and 4) a stimulus when none was present. We investigated the neutral condition as group differences were not expected to be present for this condition. We ran a factorial ANOVA with group (controls, synesthetes, schizophrenia patients) as a factor, and age and gender as covariates. There was no effect of group ($F(2,56)=3.12$, $p=.052$, $\eta_p^2=.10$) ruling out differences in criterion across the populations as a source of the results reported in the main manuscript.
